# Genomic Insights into Ceftazidime Resistance in *Burkholderia pseudomallei*: Discovery of A172T Mutation, and Palindromic GC-Rich Repeat Sequences Facilitating *penA* Duplication and Amplification

**DOI:** 10.1101/2025.02.11.637714

**Authors:** Apichai Tuanyok, Chie Nakajima, Tiernan Noll, Md. Siddiqur Rahman Khan, Pacharapong Khrongsee, Charles A. Yowell, Yu-Ping Xiao, Vanaporn Wuthiekanun, Narisara Chantratita, Henry Heine, Kuttichantran Subramaniam, Yasuhiko Suzuki, Direk Limmathurotsakul, Ayalew Mergia

## Abstract

Ceftazidime (CAZ) resistance in *Burkholderia pseudomallei*, the causative agent of melioidosis, complicates treatment in endemic regions. This study identified a novel A172T mutation and other known *penA* mutations as critical contributors to CAZ resistance in a large Thai strain collection. Frequent gene duplication and amplification (GDA) of *penA*, likely driven by Palindromic GC-Rich Repeat Sequences (PGCRRS), highlights the urgent need for rapid diagnostics and optimized treatment strategies to manage this life-threatening disease effectively.

## TEXT

The growing threat of antimicrobial resistance in *B. pseudomallei* demands a deeper understanding of the mechanisms underlining ceftazidime (CAZ) resistance. In this study, we analyzed a collection of 58 strains, isolated from 24 patients who experienced treatment failures in Northeast Thailand over two decades, 1987 to 2007 (1). These strains were specifically chosen due to their documented Etest results indicating resistance to CAZ and amoxicillin-clavulanic acid (AMC), which developed during or after treatment (Table 1). To identify the genetic factors contributing to this resistance, we employed next-generation sequencing (NGS) on the Illumina MiSeq platform as previously described (2), followed by a detailed analysis of the sequencing data using the BWA-MEM algorithm (3). Genomic alignments were conducted against the reference genome of *B. pseudomallei* K96243, and the data were visualized using the Artemis genome browser (4). *De novo* assembly of each genome was also generated using SPAdes genome assembler (Galaxy Version 3.12.0+galaxy1) or BV-BRC (http://bv-brc.org). Sequencing data are available through GenBank (accession no. PRJNA1196838).

**Table 1.**
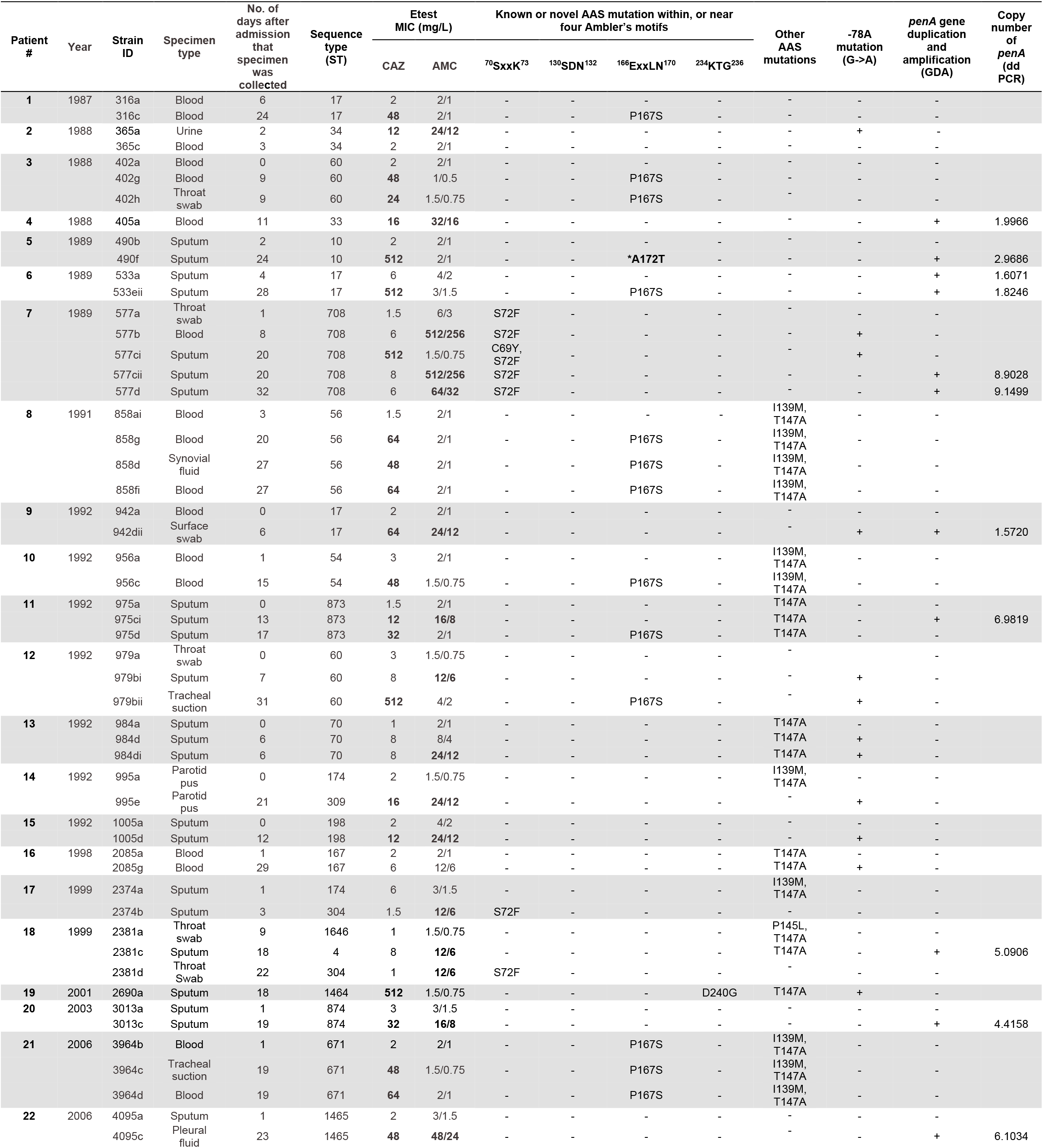
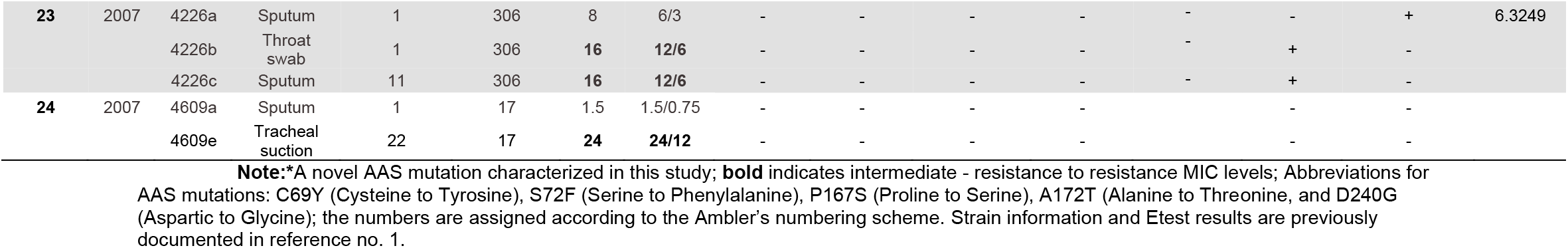
Details of *B. pseudomallei* strains used in this study, MICs, and *penA* mutations.

| Patient # | Year | Strain ID | Specimen type | No. of days after admission that specimen was collected | Sequence type (ST) | Etest MIC (mg/L) |  | Known or novel AAS mutation within, or near four Ambler's motifs |  |  |  | Other AAS mutations | -78A mutation (G→A) | penA gene duplication and amplification (GDA) | Copy number of penA (dd PCR) |
| --- | --- | --- | --- | --- | --- | --- | --- | --- | --- | --- | --- | --- | --- | --- | --- |
|  |  |  |  |  |  | CAZ | AMC | <sup>70</sup>SxxK<sup>73</sup> | <sup>130</sup>SDN<sup>132</sup> | <sup>166</sup>ExxLN<sup>170</sup> | <sup>234</sup>KTG<sup>236</sup> |  |  |  |  |
| 1 | 1987 | 316a | Blood | 6 | 17 | 2 | 2/1 | - | - | - | - | - | - | - |  |
|  |  | 316c | Blood | 24 | 17 | 48 | 2/1 | - | - | P167S | - | - | - | - |  |
| 2 | 1988 | 365a | Urine | 2 | 34 | 12 | 24/12 | - | - | - | - | - | + | - |  |
|  |  | 365c | Blood | 3 | 34 | 2 | 2/1 | - | - | - | - | - | - | - |  |
| 3 | 1988 | 402a | Blood | 0 | 60 | 2 | 2/1 | - | - | - | - | - | - | - |  |
|  |  | 402g | Blood | 9 | 60 | 48 | 1/0.5 | - | - | P167S | - | - | - | - |  |
|  |  | 402h | Throat swab | 9 | 60 | 24 | 1.5/0.75 | - | - | P167S | - | - | - | - |  |
| 4 | 1988 | 405a | Blood | 11 | 33 | 16 | 32/16 | - | - | - | - | - | - | + | 1.9966 |
| 5 | 1989 | 490b | Sputum | 2 | 10 | 2 | 2/1 | - | - | - | - | - | - | - |  |
|  |  | 490f | Sputum | 24 | 10 | 512 | 2/1 | - | - | *A172T | - | - | - | + | 2.9686 |
| 6 | 1989 | 533a | Sputum | 4 | 17 | 6 | 4/2 | - | - | - | - | - | - | + | 1.6071 |
|  |  | 533eii | Sputum | 28 | 17 | 512 | 3/1.5 | - | - | P167S | - | - | - | + | 1.8246 |
| 7 | 1989 | 577a | Throat swab | 1 | 708 | 1.5 | 6/3 | S72F | - | - | - | - | - | - |  |
|  |  | 577b | Blood | 8 | 708 | 6 | 512/256 | S72F | - | - | - | - | + | - |  |
|  |  | 577ci | Sputum | 20 | 708 | 512 | 1.5/0.75 | C69Y, S72F | - | - | - | - | + | - |  |
|  |  | 577cii | Sputum | 20 | 708 | 8 | 512/256 | S72F | - | - | - | - | - | + | 8.9028 |
|  |  | 577d | Sputum | 32 | 708 | 6 | 64/32 | S72F | - | - | - | - | - | + | 9.1499 |
| 8 | 1991 | 858ai | Blood | 3 | 56 | 1.5 | 2/1 | - | - | - | - | I139M, T147A | - | - |  |
|  |  | 858g | Blood | 20 | 56 | 64 | 2/1 | - | - | P167S | - | I139M, T147A | - | - |  |
|  |  | 858d | Synovial fluid | 27 | 56 | 48 | 2/1 | - | - | P167S | - | I139M, T147A | - | - |  |
|  |  | 858fi | Blood | 27 | 56 | 64 | 2/1 | - | - | P167S | - | I139M, T147A | - | - |  |
| 9 | 1992 | 942a | Blood | 0 | 17 | 2 | 2/1 | - | - | - | - | - | - | - |  |
|  |  | 942dii | Surface swab | 6 | 17 | 64 | 24/12 | - | - | - | - | - | + | + | 1.5720 |
| 10 | 1992 | 956a | Blood | 1 | 54 | 3 | 2/1 | - | - | - | - | I139M, T147A | - | - |  |
|  |  | 956c | Blood | 15 | 54 | 48 | 1.5/0.75 | - | - | P167S | - | I139M, T147A | - | - |  |
| 11 | 1992 | 975a | Sputum | 0 | 873 | 1.5 | 2/1 | - | - | - | - | T147A | - | - |  |
|  |  | 975ci | Sputum | 13 | 873 | 12 | 16/8 | - | - | - | - | T147A | - | + | 6.9819 |
|  |  | 975d | Sputum | 17 | 873 | 32 | 2/1 | - | - | P167S | - | T147A | - | - |  |
| 12 | 1992 | 979a | Throat swab | 0 | 60 | 3 | 1.5/0.75 | - | - | - | - | - | - | - |  |
|  |  | 979bi | Sputum | 7 | 60 | 8 | 12/6 | - | - | - | - | - | + | - |  |
|  |  | 979bii | Tracheal suction | 31 | 60 | 512 | 4/2 | - | - | P167S | - | - | + | - |  |
| 13 | 1992 | 984a | Sputum | 0 | 70 | 1 | 2/1 | - | - | - | - | T147A | - | - |  |
|  |  | 984d | Sputum | 6 | 70 | 8 | 8/4 | - | - | - | - | T147A | + | - |  |
|  |  | 984di | Sputum | 6 | 70 | 8 | 24/12 | - | - | - | - | T147A | + | - |  |
| 14 | 1992 | 995a | Parotid pus | 0 | 174 | 2 | 1.5/0.75 | - | - | - | - | I139M, T147A | - | - |  |
|  |  | 995e | Parotid pus | 21 | 309 | 16 | 24/12 | - | - | - | - | - | + | - |  |
| 15 | 1992 | 1005a | Sputum | 0 | 198 | 2 | 4/2 | - | - | - | - | - | - | - |  |
|  |  | 1005d | Sputum | 12 | 198 | 12 | 24/12 | - | - | - | - | - | + | - |  |
| 16 | 1998 | 2085a | Blood | 1 | 167 | 2 | 2/1 | - | - | - | - | T147A | - | - |  |
|  |  | 2085g | Blood | 29 | 167 | 6 | 12/6 | - | - | - | - | T147A | + | - |  |
| 17 | 1999 | 2374a | Sputum | 1 | 174 | 6 | 3/1.5 | - | - | - | - | I139M, T147A | - | - |  |
|  |  | 2374b | Sputum | 3 | 304 | 1.5 | 12/6 | S72F | - | - | - | - | - | - |  |
| 18 | 1999 | 2381a | Throat swab | 9 | 1646 | 1 | 1.5/0.75 | - | - | - | - | P145L, T147A | - | - |  |
|  |  | 2381c | Sputum | 18 | 4 | 8 | 12/6 | - | - | - | - | T147A | - | + | 5.0906 |
|  |  | 2381d | Throat Swab | 22 | 304 | 1 | 12/6 | S72F | - | - | - | - | - | - |  |
| 19 | 2001 | 2690a | Sputum | 18 | 1464 | 512 | 1.5/0.75 | - | - | - | D240G | T147A | + | - |  |
| 20 | 2003 | 3013a | Sputum | 1 | 874 | 3 | 3/1.5 | - | - | - | - | - | - | - |  |
|  |  | 3013c | Sputum | 19 | 874 | 32 | 16/8 | - | - | - | - | - | - | + | 4.4158 |
| 21 | 2006 | 3964b | Blood | 1 | 671 | 2 | 2/1 | - | - | P167S | - | I139M, T147A | - | - |  |
|  |  | 3964c | Tracheal suction | 19 | 671 | 48 | 1.5/0.75 | - | - | P167S | - | I139M, T147A | - | - |  |
|  |  | 3964d | Blood | 19 | 671 | 64 | 2/1 | - | - | P167S | - | I139M, T147A | - | - |  |
| 22 | 2006 | 4095a | Sputum | 1 | 1465 | 2 | 3/1.5 | - | - | - | - | - | - | - |  |
|  |  | 4095c | Pleural fluid | 23 | 1465 | 48 | 48/24 | - | - | - | - | - | - | + | 6.1034 |
| **23** | 2007 | 4226a | Sputum | 1 | 306 | 8 | 6/3 | - | - | - | - | - | - | + | 6.3249 |
|  |  | 4226b | Throat swab | 1 | 306 | **16** | **12/6** | - | - | - | - | - | + | - |  |
|  |  | 4226c | Sputum | 11 | 306 | **16** | **12/6** | - | - | - | - | - | + | - |  |
| **24** | 2007 | 4609a | Sputum | 1 | 17 | 1.5 | 1.5/0.75 | - | - | - | - |  | - | - |  |
|  |  | 4609e | Tracheal suction | 22 | 17 | **24** | **24/12** | - | - | - | - |  | - | - |  |

**Note:**\*A novel AAS mutation characterized in this study; **bold** indicates intermediate - resistance to resistance MIC levels; Abbreviations for AAS mutations: C69Y (Cysteine to Tyrosine), S72F (Serine to Phenylalanine), P167S (Proline to Serine), A172T (Alanine to Threonine, and D240G (Aspartic to Glycine); the numbers are assigned according to the Ambler's numbering scheme. Strain information and Etest results are previously documented in reference no. 1.

Our findings revealed a complex landscape of genetic alterations associated with CAZ and AMC resistance. We identified eight distinct amino acid substitutions (AAS) in the *penA* gene, which encodes a class A β-lactamase enzyme known to confer resistance to β-lactam antibiotics, especially CAZ and AMC (5-7) based on Etest results (Table 1). Among these substitutions, five were known to be responsible for CAZ, AMC, or imipenem (IMP) resistance (Fig. 1), while three were novel AAS mutations, I139M, P145L, and A172T. We observed that P174L, the most recently reported AAS mutation associated CAZ resistance in Hainan, China (8), was not present in our strains. Among the novel mutations, only the A172T (Alanine to Threonine) substitution near ^166^ExxLN^170^, one of the Ambler’s motifs (9) in *B. pseudomallei* strain 490f showed a particularly strong association with increased minimal inhibitory concentrations (MICs) for CAZ. This mutation was absent in the CAZ-susceptible strain 490b, which was isolated three weeks earlier from the same patient during the hospitalization. To confirm whether this mutation was responsible for the increased CAZ MIC, allelic exchange mutagenesis was performed to introduce the A172T mutant in *penA* gene of *B. pseudomallei* Bp82, a biosafe and CAZ-susceptible strain (10, 11); see Supplemental Text 1. The resulting mutant exhibited a 16-fold-increase in CAZ MIC, confirming the mutation’s critical role in resistance.

**Fig. 1.**
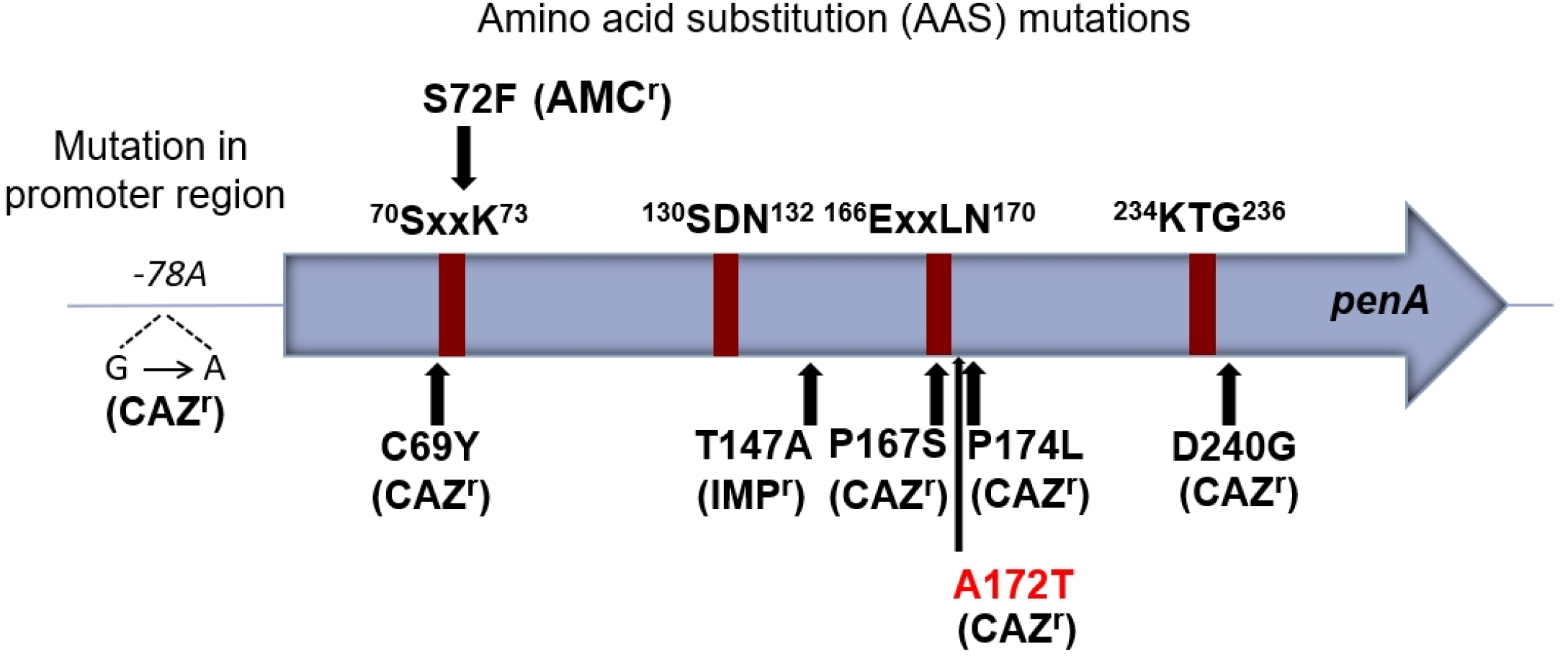
Amino acid substitution (AAS) mutations in PenA and in a promoter region known to be associated with amoxiclav resistance (AMC^r^), imipenem resistance (IMP^r^), or ceftazidime resistance (CAZ^r^). A172T is a novel AAS mutation responsible for CAZ resistance identified and characterized in this study.

In addition to AAS mutations, we identified a promoter-up mutation, specifically the -*78A* mutation (12), of the *penA* gene in the CAZ-resistant *B. pseudomallei* strains isolated from 10 (41.6%) of the 24 patients. This mutation is known to enhance the expression of *penA*, thereby increasing the production of the β-lactamase enzyme and contributing to the observed resistance. Another major resistance mechanism identified was gene duplication and amplification (GDA) of the *penA* gene, observed in 12 CAZ-resistant strains from 10 (41.7%) of the 24 patients analyzed (Fig. 2). Digital droplet PCR assays (Bio-Rad QX200™ Droplet Digital™ PCR System) using genomic DNA from *B. pseudomallei* K96243 as a single copy *penA* control confirmed that *penA* gene copies varied from two to nine among these strains, suggesting that *penA* GDA significantly increased resistance by enhancing beta-lactamase enzyme production as previously reported by us (2). In most GDA events, the junction sequences, ranging from 5 to 16 bp in length and enriched in GC content, likely resulted from homologous recombination between cruciform structures, such as stem-loop cruciform or four-way junction cruciform sequences, found at both ends of the GDA region as exemplified by strain 405a (Fig. 3). We termed this genetic structure “Palindromic GC-Rich Repeat Sequences” (PGCRRS), as it appeared to mediate the homologous recombination of the GDA. We hypothesize that these cruciform structures induce replication stress in *B. pseudomallei* under CAZ selection, leading to double-strand breaks and complicating the DNA repair processes, thereby contributing to GDA. GDA also resulted in extra copies of variable genomic regions. For example, two copies of genes BPSS0935 – BPSS1017 were found in strain 533eii, while nine copies of genes between BPSS0935 – BPSS0949 were observed in strains 577cii and 577d (Fig. 2). To confirm the GDA, initially identified in Illumina assembly contigs, we utilized the Oxford Nanopore Technologies (ONT) sequencing platform, which offers longer reads to effectively cover the GDA’s junctions. This approach confirmed the junction sequences and genomic regions affected by the GDA in three selected strains: 490f, 533a, and 942dii (BioSample accession numbers SAMN45667634, SAMN45667635, and SAMN45667647, respectively). The GDA events in all 12 strains reported in this study are described in Supplemental Table 1. The addition of ONT sequencing and the hybrid genomic assembling approach for strain 490f, as an example, are described in Supplemental Text 2.

**Fig. 2.**
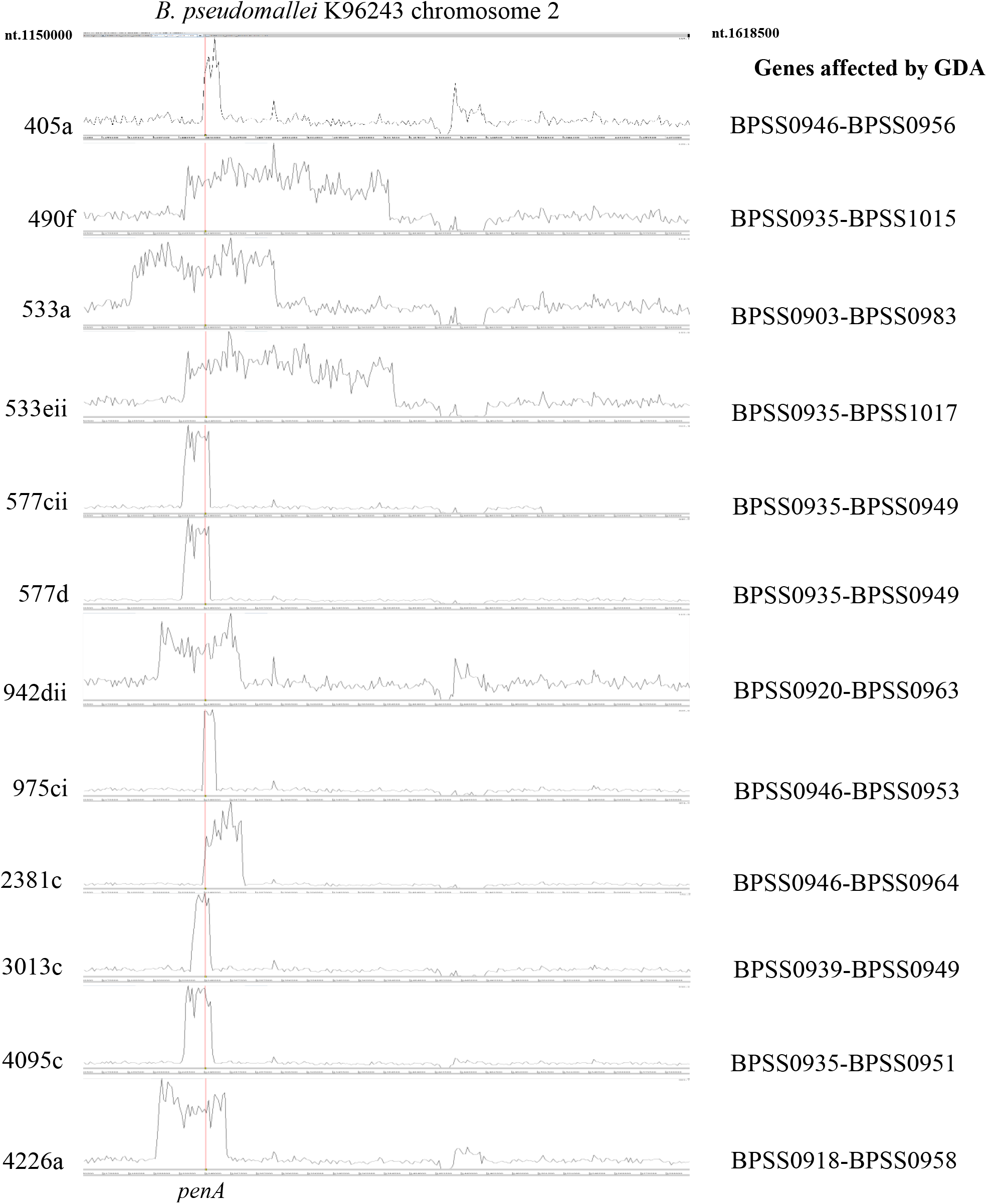
The gene duplication and amplification (GDA) of the *penA* gene, highlighting affected genomic regions observed from mapping Illumina short reads of 12 CAZ-resistant *B. pseudomallei* strains against chromosome 2 of the reference *B. pseudomallei* K96243. Amplified regions, indicated by peaks in read depth, with the *penA* highlighted in red.

**Fig. 3.**
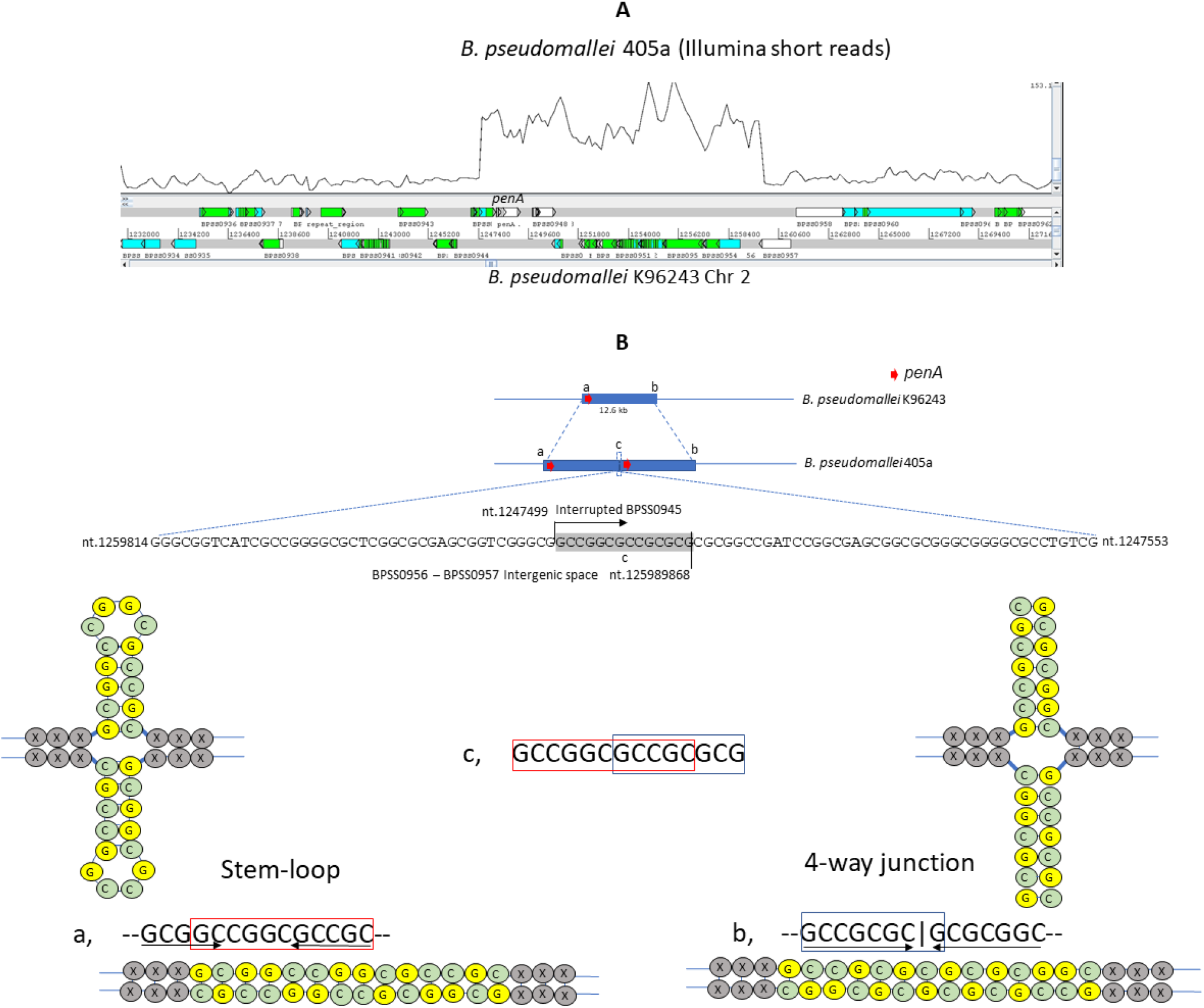
An example of gene duplication and amplification (GDA) of *penA* mediated by Palindromic GC-Rich Repeat Sequences (PGCRRS). **Panel A**: Genomic region containing *penA* and its neighboring genes, observed by mapping Illumina short reads from *B. pseudomallei* 405a against chromosome 2 of the reference strain K96243. **Panel B**: Hypothetical model illustrating GDA mediated by the recombination of the PGCRRS features, including: (a) a cruciform stem-loop structure, (b) a cruciform four-way junction structure, and (c) a 14-bp sequence resulting from the recombination between (a) and (b). **Note:** The genome coordinates shown correspond those of chromosome 2 of *B. pseudomallei* K96243.

Additionally, we have observed significantly higher MICs for CAZ or AMC in strains that contained an AAS mutation in combination with the *-78A* promoter-up mutation and/or GDA. Notable, in strain 577ci, the presence of C69Y mutation occurring against the background of the S72F mutation within the Ambler’s motif ^70^SxxK^73^ in an earlier strain 577b, shifted the resistance phenotype from AMC to CAZ. This finding suggests that modifications to PenA’s active site may influence substrate specificity. Further investigation using the artificial intelligence - driven crystal structure analysis and molecular docking approach is warranted to elucidate the underlying mechanisms. However, in this current study, we were unable to determine the genetic or molecular basis of CAZ resistance in strain 4609e from patient #24. The *penA* and other CAZ resistance - associated genes, including penicillin-binding protein 3 (*BPSS1219*) (13) in this strain were identical to those in the earlier CAZ-susceptible strain 4609a from the same patient. This observation underscores the need for additional research to identify alternative mechanisms contributing to CAZ resistance in such cases.

On another note, multi-locus sequence typing (MLST) analysis revealed that in most patients, initial isolates and resistant strains shared the same sequence types (STs), except in three patients (Table 1: patient #14, #17, and #18), the resistant strains had different STs from those initially identified at admission. This suggests that the resistant strains arose from distinct *B. pseudomallei* subpopulations that were not detected in the initial samples. Given the genetic diversity of *B. pseudomallei* in soil as previously described (14), it is possible that these patients were exposed to multiple strain genotypes. These results highlight the genetic heterogeneity of *B. pseudomallei* and the potential for multiple subpopulations to contribute to treatment failures, complicating efforts to manage CAZ resistance in clinical settings.

In conclusion, our findings provide critical insights into the genetic and molecular basis of CAZ resistance in *B. pseudomallei*. The identification of the novel A172T mutation and other previously known AAS mutations, the -*78A* promoter-up mutation, and the frequent occurrence of *penA* GDA events significantly advance our understanding of how this pathogen adapts under antibiotic pressure. Additionally, *penA*-mediated CAZ resistance appears to be increasing, as reported in multiple recent studies (8, 15). These findings from us and others emphasize the importance of developing rapid diagnostic assays that can detect these genetic alterations, guiding more effective treatment decisions in clinical practice. Implementing these insights could lead to improved management of melioidosis, ultimately reducing the morbidity and mortality associated with this challenging disease. Future research should focus on developing tools to monitor CAZ resistance in clinical settings in real-time and exploring potential therapeutic strategies that target these resistance mechanisms. Moreover, public health initiatives aimed at optimizing antibiotic use in melioidosis treatment could help mitigate the treatment failure, preserving the efficacy of CAZ and other critical antibiotics in managing this deadly disease.

## ACKNOWNLEDGEMENTS

This research was supported in part by Japan Agency for Medical Research and Development (AMED) under Grant Numbers JP24wm0125008 and JP243fa627005; CRDF Global Grant OISE-9531011; and the Wellcome Trust [220211/Z/20/Z]. A.T. and C.N. received the US-Japan Cooperative Medical Sciences Programs (USJCMSP) Collaborative Award. T.N. was supported by The Lisa Conti Florida Veterinary Scholars Fund and the Linda F Hayward Florida Veterinary Scholars Program. For the purpose of Open Access, the author has applied a CC BY public copyright license to any Author Accepted Manuscript version arising from this submission.

## Supplemental Text 1

To generate a single base A172T substitution in PenA (BP1026B_II1037), we modified a technique from López CM et al., 2009 [1]. Briefly, approximately 650 base pairs upstream and 700 base pairs downstream of the target nucleotide were amplified using the following four primers in a single PCR reaction. These primers included:

penA A172T 5’_fwd: 5’-GCTGAACACGACGCTGCCCGGCGACGAG-3’,

penA A172T 5’_rev: 5’-CGGGCAGCGTCGTGTTCAGCTCAGGCTC-3’,

penA A172T 3’_fwd: 5’-GCTGAACACGACGCTGCCCGGCGACGAG-3’, and

penA A172T 3’_rev: 5’-GGGATAACAGGGTAATCCCGATACCGGCATCGTTTCGCTGCG-3’.

The PCR fragments were then gel purified using the Zymoclean Gel DNA Recovery Kit. To construct the pExKm5-A172T PenA plasmid, the pExKm5 plasmid was digested using *EcoRI*-HF and *NotI* (New England Biolabs, NEB) and gel purified alongside the PCR products. Note: The plasmid pExKm5 was kindly provided by Dr. Schweizer at the University of Florida. The digested plasmid and PCR fragments were assembled using the NEBuilder HiFi DNA Assembly Master Mix (New England Biolabs, NEB), following the manufacturer’s protocol. The assembled plasmid was then transformed into DH5α *E. coli* on LB plates supplemented with 50 μg/ml of kanamycin (Km) and 5-bromo-4-chloro-3-indolyl-β-d-galactopyranoside (X-Gal). White colonies were selected for plasmid amplification. *B. pseudomallei* Bp82 electrocompetent cells were prepared in-house by washing three times with 300 mM sucrose, as described by Choi KH et al., 2005 [2]. Two microliters of purified plasmid were electroporated into 100 μL of competent cells using a 0.2 mm cuvette (Bio-Rad) at 2500 V, 200 Ω. The transformed cells were recovered in LB broth for 2 hours with slow shaking, then plated on LB plates containing 250 μg/mL Km and 50 μg/mL X-Gluc for 48 hours. A blue Km-resistant colony was picked, and the merodiploid was resolved on 15% sucrose YT agar supplemented with 0.8 μg/mL adenine for 48 hours. White colonies were selected on LB adenine agar supplemented with 32 μg/ml ceftazidime. The resistant colonies were then screened by PCR using penA A172T 5’_fwd and penA A172T 3’_rev primers, followed by amplicon sequencing to confirm the mutation. Minimal inhibitory concentration (MIC) of ceftazidime against a PCR positive clone (CAY2) using a test strip (Liofilchem™ MTS™ Ceftazidime [CAZ] 0.016-256 μg/mL). The MIC for the mutant strain was determined as 32 ug/mL, while Bp82 had an MIC of 1.5-2 μg/mL (see below). Whole genome sequencing by Illumina was used to confirm the presence of A172T allele in *penA* of the mutant CAY2 (BioSample# SAMN46427263).

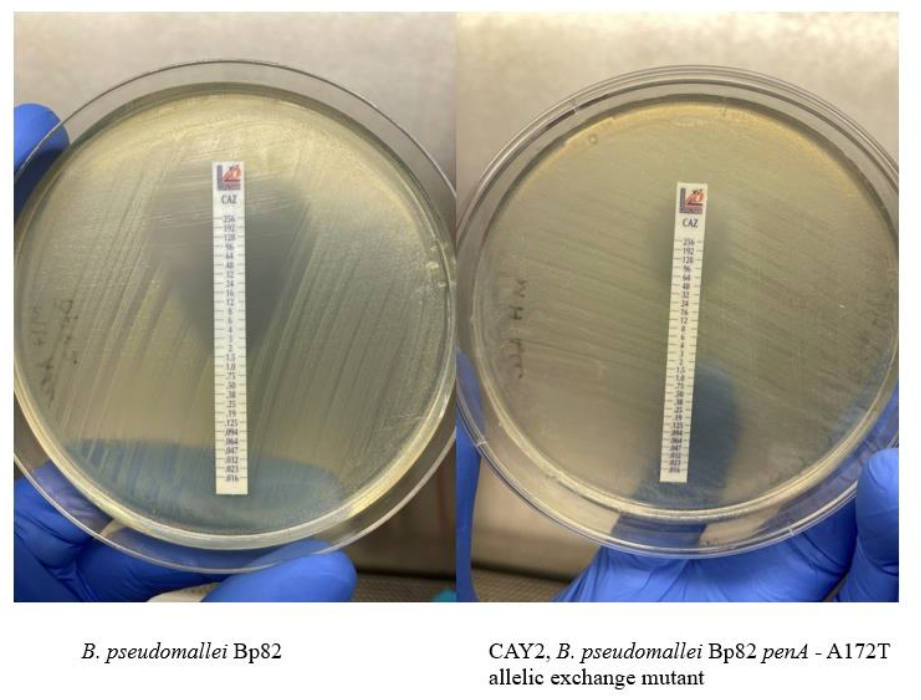

## Supplemental Text 2

The genome of *B. pseudomallei* strain 490f was sequenced using a hybrid approach combining long-read Oxford Nanopore Technologies (ONT) and short-read Illumina sequencing. Briefly, the bacteria were cultured in LB broth at 37°C with 250 rpm shaking overnight. DNA was extracted using the Promega’s Wizard® Genomic DNA Purification Kit according to the manufacturer’s instructions.

### Oxford Nanopore Sequencing

ONT sequencing was conducted on the GridION platform using an R10.4.1 flow cell and the PCR-free ONT Ligation Sequencing Kit (SQK-NBD114.24) in combination with the NEBNext® Companion Module (E7180L). The sequencing yielded 157,168 reads with an average read length of 3,165 bp. Reads were filtered for a minimum length of 2,000 bp using SeqKit v2.4.0 [1], resulting in 69,268 reads with an average read length of 5,629 bp. Genome assembly was performed with Canu v2.2 [2], producing two circular chromosome scaffolds with an average coverage of 57.46x. Unless otherwise noted, default parameters were used for all software.

### Illumina Sequencing

Illumina libraries were prepared using the Illumina DNA Prep Kit and NEBNext® Multiplex Oligos for Illumina (dual-indexed primers), targeting a 280-bp insert size. Paired-end sequencing (2 × 151 bp) was performed on the Illumina NextSeq platform, generating 18,124,202 reads. Quality control and adapter trimming were performed with bcl-convert1 v4.2.4. Reads were aligned to the ONT-derived genome scaffolds using Bowtie2 v2.4.2 [3]. Pilon v1.23 [4] was used to polish the draft assembly, resulting in a final average coverage of 749x.

### Repeat sequence correction

Initial analysis of the polished sequences in CLC Genomics Workbench v20.0.4 revealed a threefold increase in read coverage of the long reads at coordinates 2.1–2.3 Mb on chromosome 2. A 150-kb region from this area was extracted and manually re-aligned to improve mapping accuracy. To resolve this, ambiguous bases (N) were added upstream and downstream of the sequence, followed by re-mapping. There was a subset of three identical repeat sequences, two of which had unique sequence endings that aligned upstream and downstream of the 150-kb region. The three repeat regions were then internally connected each other. These adjustments ensured proper assignment of the three repeat regions.

